# Reversal of elevated *Gli3* in Autosomal Recessive Polycystic Kidney Disease does not alter cystogenesis

**DOI:** 10.1101/2024.09.18.613676

**Authors:** Lauren G Russell, Maria Kolatsi-Joannou, Laura Wilson, Jennifer C Chandler, Nuria Perretta Tejedor, Georgie Stagg, Karen L Price, Christopher J Rowan, Tessa Crompton, Norman D Rosenblum, Paul JD Winyard, David A Long

## Abstract

Polycystic kidney diseases (PKD) are genetic disorders characterised by the formation of fluid-filled cysts, which disrupt kidney architecture and function. Autosomal recessive PKD (ARPKD) is a rare form of PKD, caused by mutations in *PKHD1,* and clinically more severe than the more common autosomal dominant PKD (ADPKD). Prior studies have implicated the ciliary-located Hedgehog (Hh) pathway in ADPKD, with increased levels of Hh components in experimental ADPKD models, and reduced cystogenesis following pharmacological Hh inhibition. In contrast, the role of the Hh pathway in ARPKD is poorly understood. We hypothesised that Hh pathway activity would be elevated during ARPKD pathogenesis, and its modulation may inhibit cystogenesis, akin to prior findings in ADPKD. To test this, we utilised *Cpk* mice, a model which replicates the pathophysiology of ARPKD, and generated a human cellular ARPKD 3-dimensional cystogenesis model by mutating *PKHD1* in human collecting duct cells through CRISPR-Cas9 technology. We found significantly elevated levels of the Hh transcriptional effector *Gli3* in the *Cpk* mouse, a finding replicated in our human cellular ARPKD model. In the *Cpk* mouse, we also observed an increase in total GLI3 and GLI3 repressor protein levels. However, reduction of increased *Gli3* levels via genetic deletion in the *Cpk* mouse did not affect cyst formation. Similarly, lowering *GLI3* transcript to wildtype levels, did not influence cyst size in our human cellular ARPKD model. Collectively, these data show that elevated Gli3 does not modulate cyst progression in the context of ARPKD, highlighting the complexity of the Hh pathway in PKD.

**New and Noteworthy:** The role of the Hedgehog pathway in autosomal recessive polycystic kidney disease (ARPKD) is poorly understood. Here, we describe elevated levels of *Gli3,* the Hedgehog transcriptional effector, in murine and human ARPKD models. However, reversal of the increase in *Gli3* did not significantly affect cystogenesis in a human cell model of ARPKD or disease progression in a mouse model which replicates ARPKD pathophysiology. Collectively, our data indicates that Gli3 does not modulate ARPKD progression.

## Introduction

Polycystic kidney disease (PKD) is characterised by the formation of fluid-filled renal cysts, which disrupt the architecture and function of the kidney and is the most common genetic cause of end stage kidney disease (ESKD). The two main forms of PKD are autosomal dominant PKD (ADPKD) and autosomal recessive PKD (ARPKD) which are caused by mutations in *PKD1* or *PKD2* (encoding polycystin-1 and -2) and *PKHD1* (encoding for fibrocystin) respectively (1). ARPKD has a prevalence of 1:20,000, usually manifesting *in utero*, perinatally or during childhood, and presenting with greatly enlarged cystic kidneys. In ARPKD, early proximal tubular dilation is observed in the kidney during fetal development, which then shifts to dilation and cyst formation in the distal tubules and predominantly the collecting duct (2). Disease progression occurs rapidly leading to the presence of cysts throughout the kidney destroying the normal renal architecture, along with interstitial fibrosis (1). Patients with ARPKD frequently progress to ESKD and require renal replacement therapy. Currently the only approved treatment for adults with ADPKD is tolvaptan, a vasopressin receptor 2 antagonist targeting cAMP signalling. While there are two ongoing phase 3 trials for tolvaptan in infants and children, it is not yet approved for children with PKD and there are no other treatment options for ARPKD patients (3). Thus, there is a need for a greater understanding of the mechanisms of disease in ARPKD to aid the development of novel treatments.

The disease-causing genes mutated in PKD patients localise to the primary cilia. This has led to an interest in the ciliary-located Hedgehog (Hh) pathway as a possible mediator in the pathogenesis of PKD (4). Hh signalling is activated through the binding of Hh ligands to the transmembrane receptor, Patched1 (PTCH1). In the absence of Hh ligands, smoothened (SMO), a G protein-coupled receptor, is constitutively inhibited by PTCH1. This prevents the translocation of SMO to the primary cilium which allows for the phosphorylation of the GLI transcription factors (GLI1, GLI2 and GLI3). Phosphorylated GLI proteins are partially degraded by the proteosome to form a truncated repressive form of GLI, which translocates to the nucleus and represses the transcription of Hh target genes (5). When Hh ligands bind to PTCH1, the constitutive repression of SMO is blocked, leading to its translocation and accumulation in the primary cilium. GLI transcription factors are then transported to the ciliary tip in a complex with Suppressor of Fused (SUFU). Full-length GLI proteins dissociate from this complex and translocate to the nucleus to activate the transcription of genes, such as those that regulate the cell cycle, proliferation and apoptosis (5, 6).

Components of the Hh signalling pathway are elevated in human ADPKD tissue (7, 8), mice with mutations in the ciliary genes, *Ift140*, *Thm1* and *Arl13b* (9–11); two ADPKD models, the *jck* mutant and the conditional deletion of *Pkd1* in postnatal mice (10); as well as in cystic murine metanephric kidney explants (12). Modulation of the Hh pathway through pharmacological inhibition has been demonstrated to be effective in reducing cyst progression in *in vitro* (8, 12) and mouse models (13, 14). The *Pck* ARPKD rat model also shows elevated SMO and GLI proteins in the lining of the cyst epithelium and responded to treatment with the SMO inhibitor, cyclopamine, leading to reduced renal cyst area (13). Furthermore, treatment with GANT61, a GLI1/2 inhibitor, alleviated the cystic phenotype and interstitial fibrosis in a ciliary mutant mouse (11, 14). Conversely, in a *Pkd1-*mutant mouse, the genetic overactivation or deletion of *Smo,* or deletion of both *Gli2* and *Gli3,* specifically in renal epithelial cells did not alter cyst formation. This study suggested that Hh signalling is not critical within the tubule epithelium in mediating cystogenesis caused by mutations in *Pkd1* (15). Thus, the function of Hh signalling in various PKD models is complex and remains only partially deciphered. In the context of ARPKD, Hh signalling has only been investigated in the *Pck* rat model (13), with no prior studies examining the pathway in either *in vitro* or mouse models.

In this study, we examined the role of Hedgehog signalling in both a murine and human cellular model of ARPKD. We observed significant upregulation of the downstream Hh pathway effector *Gli3* in *Cpk^-/-^* mice, a model which replicates the pathophysiology of ARPKD, and *PKHD1*-mutant human collecting duct cells. We then reversed the increased levels of Gli3 in both mouse and cellular ARPKD models and saw no effect on either cyst formation *in vitro* or disease progression *in vivo*. These results suggest that although *Gli3* is upregulated during cyst progression in ARPKD models, its modulation does not alter the cystic phenotype in ARPKD.

## Methods

### Experimental Animal and Procedures

*Cpk^+/-^* mice (16) were bred to generate wild-type and homozygous littermates for analysis. To decrease *Gli3* levels, heterozygous *Gli3^XtJ^* mice (17) were crossed with *Cpk* heterozygous mice to generate *Gli3^XtJ/+^;Cpk^+/-^* double heterozygous mice. Male or female *Gli3^XtJ/+^;Cpk^+/-^* double heterozygous mice were then crossed with either *Cpk^+/-^* and the resulting litters were collected. All lines were maintained on a C57BL/6 background. Mice of both sexes were used for analysis. Kidney and body weight were measured, and blood was collected via cardiac puncture. All procedures were approved by the UK Home Office. Blood urea nitrogen (BUN) levels were quantified in plasma using the QuantiChrom^TM^ Urea Assay Kit (BioAssay Systems).

### Histological Analysis

Histological analyses were performed on 5μm periodic acid-Schiff stained sections. Slides were imaged using the Hamamatsu NanoZoomer slide scanner at 20x magnification. All analysis was conducted blinded on FIJI ImageJ software. The average area of individual cysts was determined using Image J particle analysis tool, with cysts defined as those >10,000μm^2^ in cross-sectional area (18). The cystic index (percentage of total cyst area/total kidney area), average cyst size and number of cysts were then quantified.

### Generation of PKHD1 mutant human collecting duct cell lines by CRISPR/Cas9

Human collecting duct (HCD) cells were provided by Professor Pierre Ronco, Hopital Tenon, Paris, France, and originally derived from non-tumorous human kidney cortex, immortalised with SV-40 virus with clones selected that stained specifically for mAb272, a marker of collecting duct principal cells (19–21). HCD cells were maintained as previously described (19). Exon 5 of *PKHD1* was identified in the Leiden Open Variation Database (22) as a region to contain a high number of pathogenic variants in ARPKD patients. Therefore, a sgRNA was designed for exon 5 of *PKHD1* (5’-3’ ACTTCCTGGAAGCATACTTC) and cloned into a pX330 plasmid (Addgene plasmid, 42230). HCD cells were transfected with the sgRNA encoding plasmid and pAcGFP1-C1 plasmid using FuGENE (Promega). After 48 hours, fluorescence activated cell sorting (FACS) was performed to isolate GFP-positive cells and these were cultured for 2 weeks. Single cells were then seeded into 96-well plates by FACS, and clonal cell lines were expanded. Clones were genotyped by PCR of the targeted *PKHD1* genomic sequence and Sanger sequencing, using the following primers: 5’-3’ ACTGCTGGGAATCCTTGGTT and AGACACGCTGGCTCATTTACA. One *PKHD1*-mutant clonal cell line was generated, in addition to an isogenic wildtype control cell line that had undergone CRISPR transfection and cell sorting but no mutation in *PKHD1* was present.

### 3D cyst assay

7.3μl of Matrigel (Corning) were mixed with 2.7μl of 2000 cells in media on ice to a total volume of 10μl which was added to the inner chamber of 15-well 3D μ-slides (ibidi) and left to set at 37°C for 30-40 minutes. 50μl of media was then added per well and changed every 2 days. At day 6, the cells were imaged on an Invitrogen EVOS M5000 Microscope at 20x magnification. Fifteen images were taken per well and 3 wells were imaged per condition or cell type. Experiments were repeated independently at least 3 times, 50 cysts were quantified per experiment and a total number of 150-200 cysts per condition were measured.

### Drug treatment

For inhibition of Hedgehog signalling, cells were treated with the SMO inhibitor Cyclopamine (23). After cells were plated in Matrigel, the media added was supplemented with 10μm Cyclopamine (LKT Labs) and changed every 2 days.

### siRNA transfection

To lower *GLI3* levels *in vitro*, siRNA inhibition was used. The following conditions were used for each cell type: non-transfected, non-targeting siRNA and *GLI3* siRNA. The following siRNAs were utilised: ON-TARGETplus non-targeting siRNA Control Pool (Dharmacon, D-001810-10-5) and ON-TARGETplus Human *GLI3* siRNA SMARTPool (Dharmacon, L-011043-00-0005). 30nM of siRNA was transfected into HCD cells using Lipofectamine RNAiMAX (Invitrogen) and cells were incubated at 37°C for 6 hours. After 6 hours, cells were prepared for the 3D cyst assay or plated in 2D for RNA extraction after 6 days in culture.

### Quantitative RT-PCR

The RNeasy Plus Mini kit and RNeasy Micro Kit (Qiagen) were used for mRNA extraction from murine tissue and HCD cells, respectively. cDNA was prepared using the iScript gDNA Clear cDNA Synthesis kit (BioRad) as per the manufacturer’s instructions. Quantitative real-time polymerase chain reaction (qRT-PCR) was performed in duplicate using the qPCRBIO SyGreen Mix Lo-ROX (PCRBioSystems) with gene specific primers (*Ihh, Shh, Ptch1, Smo, Sufu, Gli1, Gli2, Gli3/GLI3*) normalised to the housekeeping gene (*Gapdh/GAPDH*).

### Western blotting

Protein lysates were prepared from 14-day postnatal kidneys by homogenisation in radioimmunoprecipitation assay (RIPA) buffer with 40μl/ml cOmplete^TM^ EDTA-free Protease Inhibitor Cocktail and 10μl/ml sodium orthovanadate phosphatase inhibitor. 50μg of protein lysates was run on a 7.5% Mini-PROTEAN TGX Precast gel (Bio-Rad) and transferred to polyvinylidene fluoride (PDVF) membranes. Membranes were blocked in 5% milk, 0.1% bovine serum albumin (BSA) in 0.1% Tween-20 in PBS and incubated in the following antibodies overnight at 4°C: Goat anti-GLI3 (1μg/ml, R&D Systems, #AF3690) or Rabbit anti-alpha-tubulin (1μg/ml, Abcam, ab4074). Membranes were then incubated in the appropriate anti-goat (DAKO, P044901) or anti-rabbit (DAKO, P044801) horseradish peroxidase (HRP)-conjugated secondary antibodies (1:2000) for 1 hour at room temperature and proteins were visualised by chemiluminescent imaging. Densitometry of the 190kDa full-length GLI3 (GLI3A) and 83kDa truncated GLI3 (GLI3R) protein bands were calculated relative to α-tubulin on FIJI ImageJ software. Total GLI3 was calculated as total GLI3A and GLI3R protein and the ratio of GLI3A:GLI3R was quantified.

### Statistical Analysis

All statistical analyses were performed using GraphPad PRISM. All data was plotted as mean value ± standard error of the mean (SEM). Normality was assessed by Shapiro-Wilk test and normally distributed data was analysed by an Unpaired t-test for comparisons between two groups and a One-way ANOVA with Tukey’s multiple comparisons test for comparisons with more than two groups. For data not normally distributed, data was analysed using a Mann-Whitney U-test for comparisons between two groups and Kruskal-Wallis test with Dunn’s multiple comparison test for comparisons with more than two groups. For *in vitro* experiments with two variables, such as cell type and treatment group, a two-way ANOVA with Tukey’s multiple comparison test was utilised.

## Results

### *Gli3* levels are elevated in the cystic kidneys of the *Cpk* model

Firstly, we examined the levels of components of the Hh signalling pathway in the *Cpk* mouse model, which replicates the pathophysiology of ARPKD (24–26). In this model, cystogenesis initiates during embryogenic development with the presence of tubular dilations which rapidly progress to renal cysts in the collecting ducts from birth through to postnatal day 21, resembling the phenotype of rapid cyst development seen in the human disease (25). We measured the transcript levels of Hh pathway genes in whole kidneys from *Cpk^-/-^* mice and their wildtype littermates at postnatal (P) 10, P14 and P21, when cysts are present within the tissue (**Figure 1A**). No significant differences were detected at any timepoint in the levels of either of the Hh ligands known to be expressed in the kidney *Ihh* and *Shh* (27) or the downstream Hh pathway genes, *Ptch1*, *Smo*, *Sufu, Gli1* or *Gli2* between *Cpk^+/+^* and *Cpk^-/-^* mice (**Figure 1B-D**). Notably, the transcription factor *Gli3* was significantly upregulated in *Cpk^-/-^* mice at both P14 by 2.7-fold and P21 by 3.5-fold in comparison with *Cpk^+/+^* mice (**Figure 1C, D,** P14 *p=0.005,* P21 *p=0.002*). Prior to this, at P10 there is a slight increase in *Gli3* transcript levels with a 1.3-fold change, although this is not significantly different at this timepoint (**Figure 1B**, *p=0.330*). The upregulation of *Gli3* transcript between P10 to P14 is in line with the period of rapid cyst progression observed in this model.

**Figure 1.**
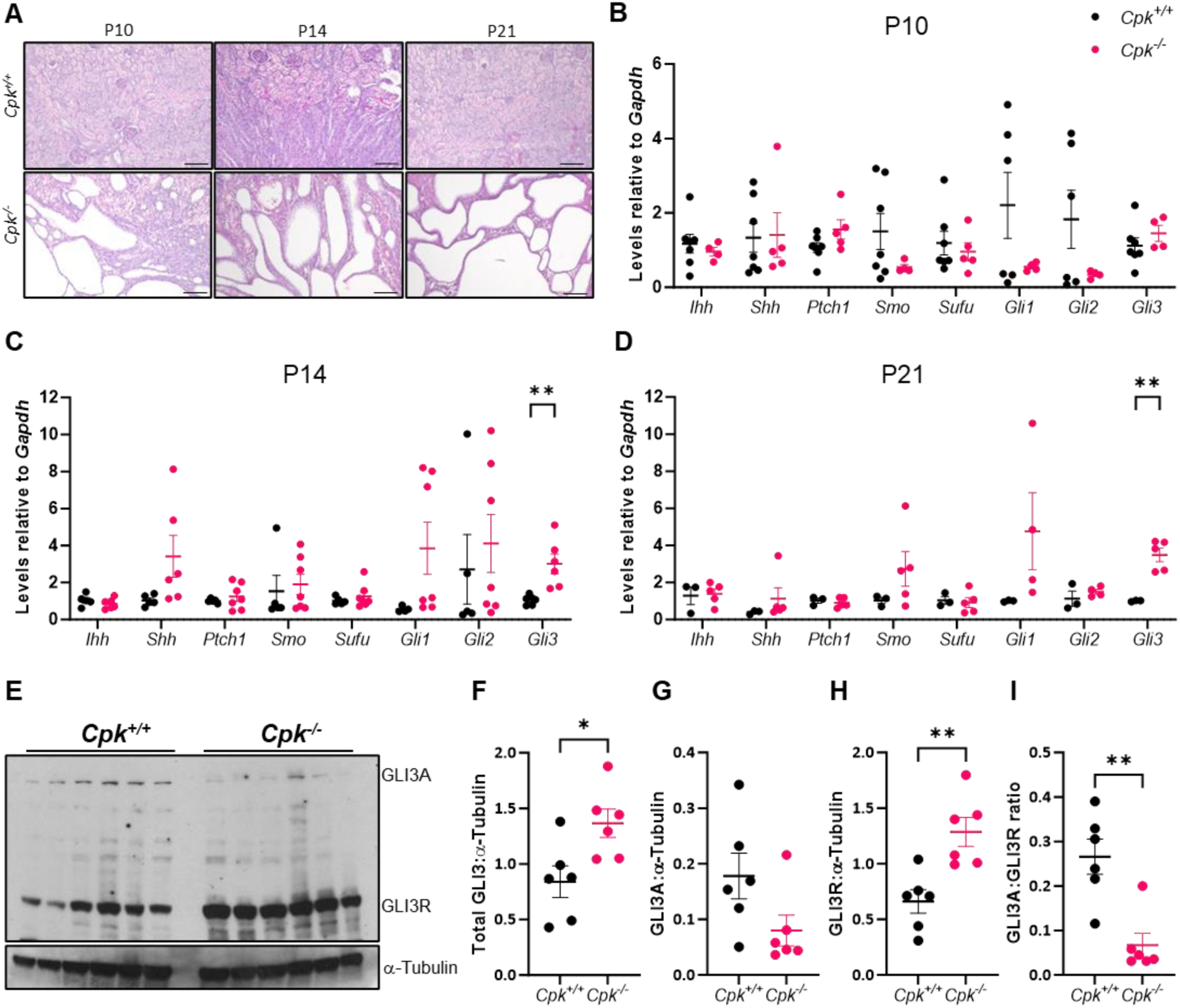
*Gli3* is elevated at both the transcript and protein level in the cystic kidneys of *Cpk^-/-^* mice. (**A**) Representative images of periodic acid-Schiff (PAS) staining on 5μm sections of *Cpk^+/+^* and *Cpk^-/-^*kidneys at P10, P14 and P21. Images taken at 10x magnification. Scale bars: 100μm. Quantitative RT-PCR analysis of *Ihh, Shh, Ptch1, Smo, Sufu, Gli1, Gli2* and *Gli3* transcript levels relative to *Gapdh* from whole kidney RNA from *Cpk^+/+^* and *Cpk^-/-^* mice at (**B**) P10, (**C**) P14 and (**D**) P21. Ct values of each gene of interest were normalised to littermate controls at each timepoint (Unpaired t-test, *Gli3* P14 *p=0.005, Gli3* P21 *p=0.002, n=7,5 3* for *Cpk^+/+^*, *n=4,6,5* for *Cpk^-/-^* mice at P10, P14 and P21 respectively). (**E**) Western blot analysis of the levels of full-length GLI3 activator (GLI3A) and truncated GLI3 repressor (GLI3R) in P14 kidney lysates from *Cpk^+/+^* (n=6) and *Cpk^-/-^* (n=6). (**F**) The relative intensity of total GLI3, a sum of GLI3A and GLI3R (*p=0.0152*), (**G**) GLI3A and (**H**) GLI3R (*p=0.009*) were quantified by densitometry in arbitrary units relative to endogenous alpha-tubulin. (**I**) The ratio of GLI3A: GLI3R was quantified (*p=0.004*) (Mann-Whitney t-test, *n=6* mice per group). Data represents mean ± SEM.

The balance between full-length activator GLI3A and repressor GLI3R protein is critical in determining the cellular function of Hh signalling (28). Therefore, the protein levels of GLI3A and GLI3R were analysed by western blotting in 14-day old *Cpk^-/-^* and wild-type littermates. Total GLI3 protein levels, a sum of both GLI3A and GLI3R protein, and GLI3R protein levels were significantly increased in *Cpk^-/-^* mice (**Figure 1E, F,** *p=0.0152* and *p=0.009* respectively). Although, there is a trend towards reduced GLI3 activator protein levels in *Cpk^-/-^* mice compared with wildtype controls, this was not significant (**Figure 1H**, *p=0.065*). There was a significant decrease in the ratio of GLI3 activator to GLI3 repressor in *Cpk^-/-^* mice, further demonstrating an increase in the relative levels of GLI3 repressor (**Figure 1I**, *p=0.004*). Collectively, these data suggest there is enhanced transcript levels of *Gli3*, alongside an increase in the total level of GLI3 protein. Additionally, a shift towards the processing of GLI3 full length activator protein to the truncated repressive form is observed in cystic kidneys in the *Cpk* model of ARPKD.

### Reduction of *Gli3* does not alter the cystic phenotype in *Cpk^-/-^* mice

We subsequently hypothesised that elevated Gli3 may be driving cystogenesis in ARPKD and that preventing this increase could reduce disease severity. To test this, we modulated *Gli3* levels genetically in the *Cpk* model using mice harbouring a *Gli3^XtJ^* allele, which contains a loss-of-function *Gli3* mutation (17). As the *Gli3^XtJ/XtJ^* mouse is embryonic lethal (17), we reduced *Gli3* levels in the *Cpk* mouse using a heterozygous *Gli3^+/XtJ^* haploinsufficiency model. We crossed *Gli3^+/XtJ^;Cpk^+/-^* double heterozygous mice with *Cpk^+/-^* mice and collected *Gli3^+/XtJ^;Cpk^-/-^* offspring and littermates at P14 (**Figure 2A**). The loss of one *Gli3* allele was sufficient to significantly reduce *Gli3* transcript levels in *Gli3^+/XtJ^;Cpk^-/-^* mice by 69% relative to *Cpk^-/-^* littermates (**Figure 2B**, *p<0.001*). However, no changes were observed in the kidney:body weight ratio or BUN levels between *Cpk^-/-^* and *Gli3^+/XtJ^;Cpk^-/-^* mice (**Figure 2C, D**). Analysis of the cystic phenotype of these mice showed no differences in cystic index between *Cpk^-/-^* and *Gli3^+/XtJ^;Cpk^-/-^* kidneys, with a mean value of 64.5±1.62% and 69.9±1.72% respectively (**Figure 2A, E**). Additionally, no difference was observed in the average cyst size or number of cysts between *Gli3^+/XtJ^;Cpk^-/-^* mice and *Cpk^-/-^* mice (**Figure 2F, G**). These data suggest that although *Gli3* is elevated in the cystic kidneys of *Cpk* mice, decreasing the level of *Gli3* does not alter the ARPKD cystic phenotype of the *Cpk* mouse.

**Figure 2.**
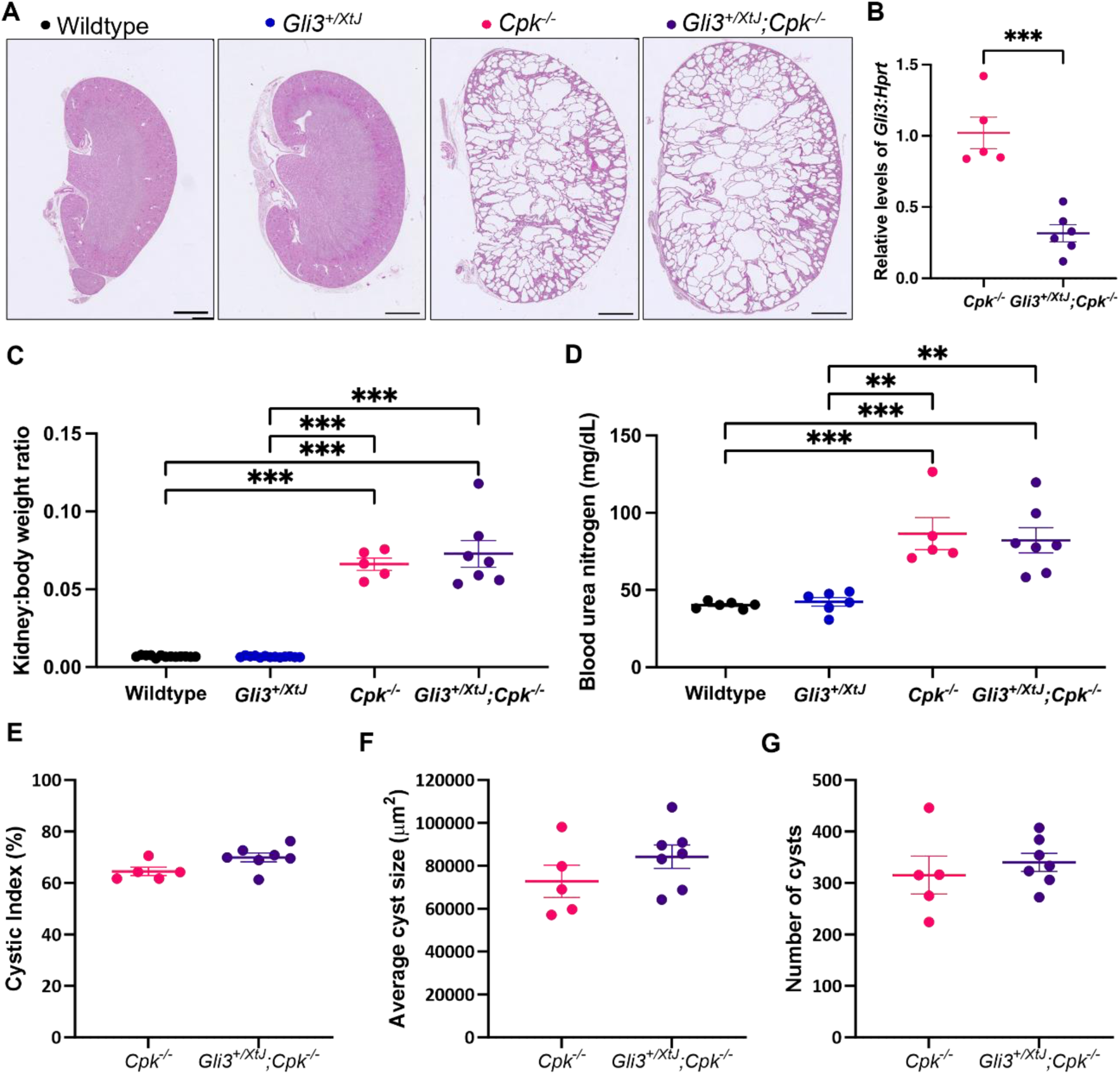
Reduced levels of *Gli3* have no effect on cyst progression in an autosomal recessive polycystic kidney disease mouse model. (**A**) Representative images of periodic acid-Schiff (PAS) staining on 5μm sections of wildtype, *Gli3^+/XtJ^, Cpk^-/-^* and *Gli3^+/XtJ^;Cpk^-/-^* kidneys at P14. 10x magnification, scale bars: 1mm. (**B**) Quantitative RT-PCR analysis of *Gli3* relative to *Hprt* from whole kidney RNA at P14. Relative Ct values of each gene of interest were normalised to *Cpk^-/-^* littermates (Unpaired t-test, *p<0.001*). (**C**) Kidney:body weight ratio and (**D**) blood urea nitrogen (BUN) levels for each group. One-way ANOVA with Tukey’s multiple comparisons test. Quantification of PAS staining was performed on *Cpk^-/-^*and *Gli3^+/XtJ^;Cpk^-/-^* mice to determine (**E**) Cystic Index, the percentage of total cyst area/ total kidney area, (**F**) average cyst size (μm2) and (**G**) total number of cysts per kidney section. Cysts were defined as ≥10,000μm^2^. Data represents mean ± SEM. Unpaired t-test. Data represents mean ± SEM. **P<0.01. n= 6 for wildtype, n=6 for *Gli3^+/XtJ^*, n=5 for *Cpk^-/-^*, and n=7 for *Gli3^+/XtJ^;Cpk^-/-^* mice.

### Generation of a 3-dimensional human cystogenesis *in vitro* model of ARPKD

To further investigate the possible role of *GLI3* in the pathogenesis of ARPKD, we generated a *PKHD1-*mutant HCD cell line using CRISPR-Cas9. We utilised CRISPR-Cas9 to target exon 5 of *PKHD1*, a region identified to have a high number of pathogenic variants reported in ARPKD patients in the Leiden Open Variation Database (22). We generated an isogenic clone with no mutations in the *PKHD1* gene following transfection and a clone with a compound heterozygous mutation in *PKHD1*, herein defined as *PKHD1-*mutant, with a single nucleotide deletion in one allele and 19 nucleotide deletion in the other allele (**Figure 3A**). Both frameshift deletions are predicted to result in a premature stop codon and result in a truncated *PKHD1* protein. We analysed the effect of this *PKHD1* mutation on *in vitro* cyst formation. Cells from both lines were embedded in Matrigel and cultured for 6 days until 3D cyst-like structures had formed. We observed significantly larger cyst structures in the *PKHD1*-mutant HCD line, with a greater average cyst area (3349±234μm^2^) when compared with the isogenic control HCD cell line (2564±110μm^2^) (**Figure 3B, C,** *p=0.016*). Additionally, the *PKHD1*-mutant HCD cell line form significantly larger individual cysts than isogenic control cells, with the largest *PKHD1*-mutant cysts measured at >8000μm^2^ compared with a maximum cyst size of 7250μm^2^ for isogenic control cysts (**Figure 3D***, p<0.001*). We found significantly elevated *GLI3* transcript levels in *PKHD1*-mutant HCD cells, increased by 2-fold, as compared with isogenic control HCD cells (**Figure 3E**, *p=0.008*). This mirrors the findings observed in the *Cpk* mouse model of ARPKD, where increased levels of *Gli3* transcript levels were detected in cystic mice. Thus, increased levels of *GLI3* transcript are common to both *in vivo* and *in vitro* models of murine and human ARPKD.

**Figure 3.**
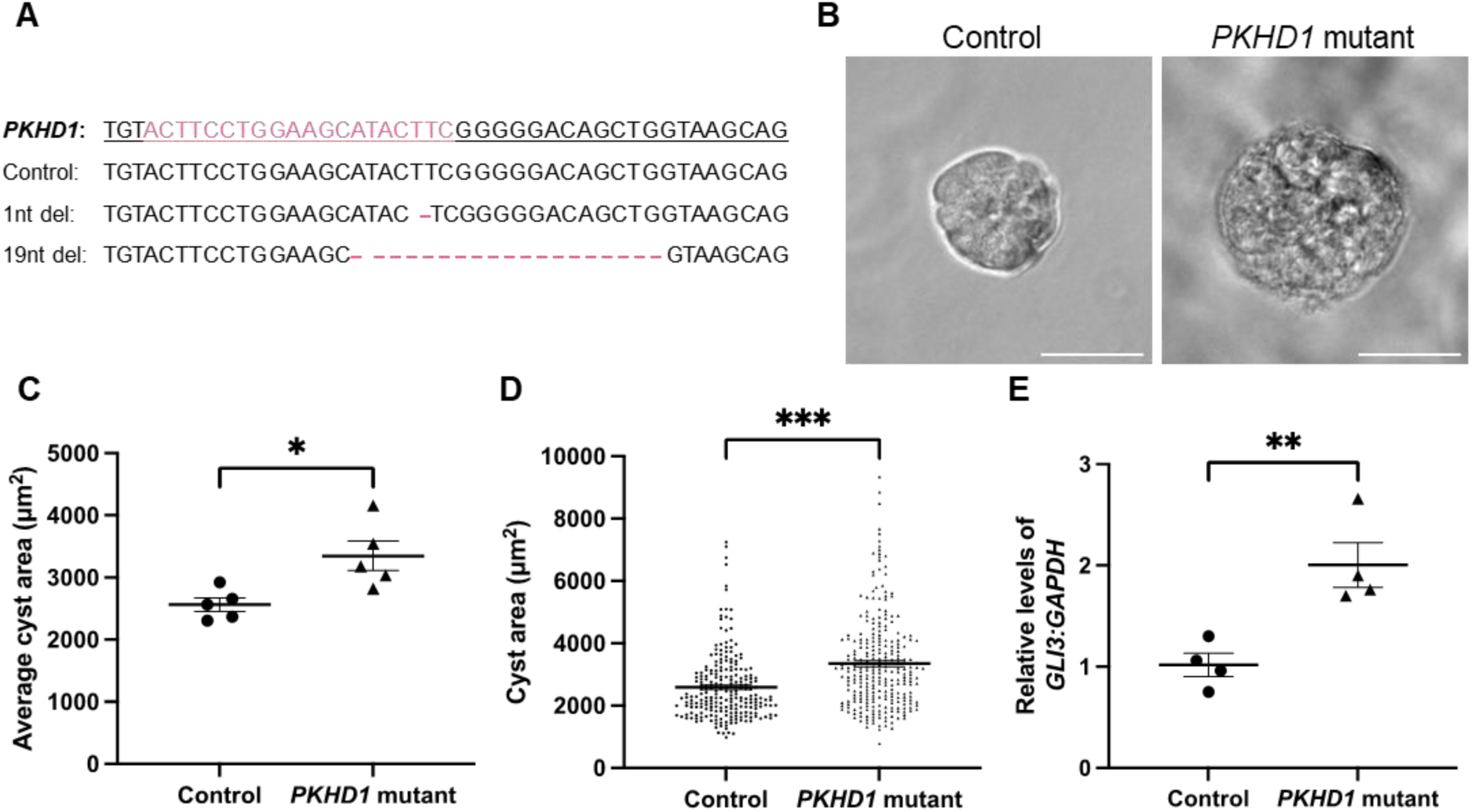
Generation of a human *in vitro* model of autosomal recessive polycystic kidney disease. (**A**) CRISPR-Cas9 was utilised to generate mutations within exon 5 of *PKHD1* in a human collecting duct cell line. Alignment of Sanger sequencing data of *PKHD1* exon 5 from clonal cell lines generated was analysed using Synthego ICE analysis. *PKHD1* wildtype sequence is underlined and sgRNA binding site is highlighted in pink. Nucleotide deletion sites are indicated with a dash. (**B**) Representative images of isogenic wildtype control and *PKHD1*-mutant 3D cyst-like structures formed in Matrigel following 6 days in culture. (**C**) The cross-sectional area of each cyst was quantified, and the average cyst area was calculated for each independent repeat. (**D**) Cyst area of each individual cyst quantified across all repeats. n=5 independent repeats. (**E**) Quantitative RT-PCR analysis of *GLI3* relative to *GAPDH* from RNA extracted from isogenic wildtype control and *PKHD1*-mutant HCD cells cultured for 6 days. Relative Ct values of each gene of interest were normalised to isogenic wildtype control HCD cells. Data represents mean ± SEM. Unpaired t-test. n=4 independent repeats. *P<0.05, **P<0.01, ***P<0.001.

### Reduction of *GLI3* does not reduce cyst formation *in vitro* in a *PKHD1-*mutant cell line

We next aimed to analyse the effect of Hh pathway inhibition on cyst formation in this cellular ARPKD model. Initially, we inhibited Hh pathway activity using the SMO inhibitor, cyclopamine, in both isogenic control and *PKHD1-*mutant HCD cells. Treatment with 10μM cyclopamine for 6 days significantly reduced average cyst size in both isogenic control and *PKHD1-*mutant cells in comparison to untreated cells (**Supplementary Figure 1**). We subsequently examined the effect of modulating *GLI3* transcript levels on cyst formation in both isogenic control and *PKHD1* mutant HCD cells. To reduce the levels of *GLI3* transcript, HCD cell lines were transfected with *GLI3* siRNA prior to plating in Matrigel. Cells were either left untransfected or transfected with 30nM of either non-targeting siRNA or *GLI3* siRNA. Transfection with the *GLI3* siRNA significantly reduced *GLI3* transcript levels by 0.55-fold in *PKHD1-*mutant HCD cells, down to similar levels observed in control HCD cells (**Figure 4A**, *p=0.016*). In isogenic control cells, *GLI3* transcript levels were reduced by an average of 0.61-fold, however the level of knockdown was variable and not significantly different (**Figure 4A**). Although *GLI3* siRNA reduced *GLI3* transcript levels in *PKHD1*-mutant cells, it did not influence cyst area with no significant difference in average cyst area between all experimental conditions for the *PKHD1*-mutant cell line (**Figure 4B, C**). There was also no significant difference in cyst area between any of the conditions for the isogenic control cell line (**Figure 4B, C**).

**Figure 4.**
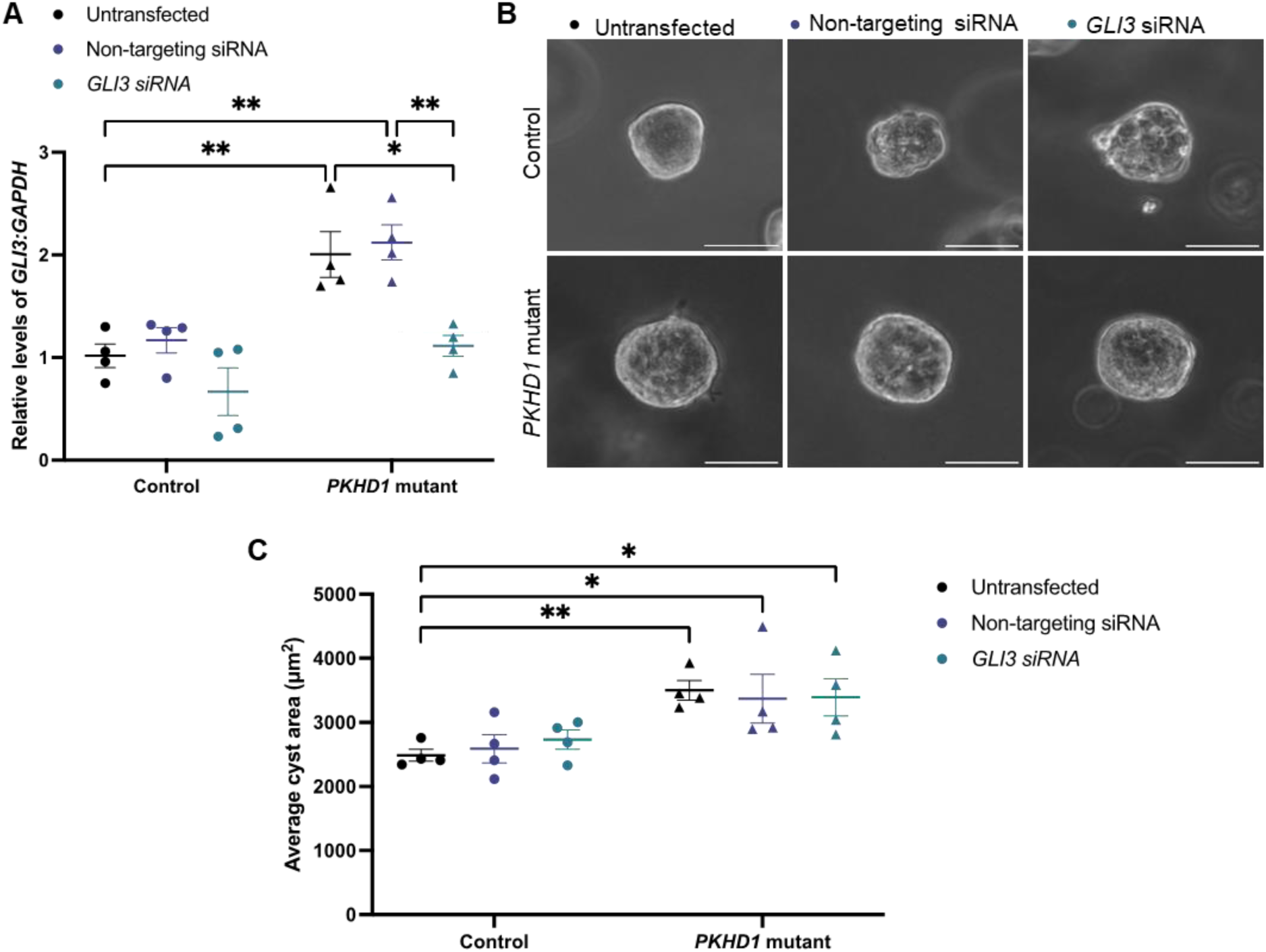
*GLI3* knockdown does not alter cyst formation *in vitro* in a human cellular ARPKD model. Isogenic wildtype control and *PKHD1*-mutant cells were transfected with non-targeting siRNA or *GLI3* siRNA and embedded in Matrigel to generate cyst structures. (**A**) Quantitative RT-PCR analysis of *GLI3* relative to *GAPDH* from RNA extracted from control and *PKHD1*-mutant HCD cells for all conditions 6 days after transfection. Relative Ct values of each gene of interest were normalised to untreated isogenic control HCD cells. (**B**) Representative images of isogenic control and *PKHD1*-mutant 3D cyst-like structures that were not transfected or transfected with non-targeting siRNA or *GLI3* siRNA. Images of the cysts were taken at day 6 at 20x magnification. (**C**) The cross-sectional area of each cyst was quantified, and the average cyst area was calculated for each independent repeat. (**D**) Cyst area of each individual cyst quantified across all repeats. n=4 independent repeats. Data represents mean ± SEM. Two-way ANOVA with Tukey’s multiple comparisons test. Statistical comparisons are shown relative to the untransfected isogenic wildtype control and between conditions of the same cell type. *P<0.05, **P<0.01.

Thus, we have demonstrated that Gli3 is significantly upregulated in two different models of ARPKD: the *Cpk* mouse and a human *PKHD1*-mutant clonal collecting duct cell line. Additionally, we have demonstrated that reducing the transcript levels of *Gli3/GLI3* levels in both the mouse and human cellular model of ARPKD does not alter cyst formation, suggesting it is not required for the cystic phenotype in ARPKD.

## Discussion

Despite studies observing an increase in Hh pathway levels in both *in vitro* and *in vivo* models of ADPKD, as well as in human ADPKD tissue (9, 10, 12) there has been a gap in research regarding the role of Hh in murine and human ARPKD. Our findings have bridged this gap by identifying a significant upregulation of the Hh transcriptional effector *Gli3* in the *Cpk* mouse. Elevation of *Gli3* transcript was in line with the period of rapid cyst progression at the later stages of the disease at P14 to P21. This provides the first evidence of *Gli3* upregulation in an ARPKD murine model, paralleling data in ADPKD models and renal cystic mouse models caused by mutations in ciliary genes (9–11). Similar to our findings, upregulation of the Hh pathway in other cystic mouse models, including *Gli3*, has only been observed at the more advanced stages of the disease (9, 11). In this study we also detected *GLI3* upregulation in a newly developed human *PKHD1*-mutant cellular model, mimicking that observed in the *Cpk-*mutant mouse.

Further to this, we found an increase in the total GLI3 protein level in *Cpk* mice at P14. Within the cell, GLI3 can act as either a transcriptional activator (GLI3A) or transcriptional repressor (GLI3R) dependent on the activity of the Hh pathway in a context dependent manner (28). In the *Cpk* model, we detected increased processing of GLI3A to GLI3R, with a tendency for reduced levels of GLI3A and elevated levels of GLI3R, indicating Hh pathway inhibition in the highly cystic kidneys of *Cpk* mice. The importance of elevated GLI3R was not explored in this study, but previous work has shown that this can lead to severe pathological effects in the kidney during its development. In mice, the global overexpression of GLI3R causes severe renal developmental abnormalities, with mice displaying hydroureter, hydronephrosis and some mutants having a bi-lobed kidney with a single ureter (29). Pallister-Hall syndrome (PHS) is associated with truncating mutations in *GLI3*, which leads to the production of a constitutively active GLI3R-like protein (30, 31). Around 27% of PHS patients have renal developmental abnormalities, most frequently these patients present with phenotypes such as hypoplasia or agenesis (31).

A common feature of PKD pathogenesis is an increase in cAMP levels, leading to the activation of kinases such as PKA or GSK3β, key regulators of GLI3 processing to GLI3R (1, 5). It could be hypothesised that enhanced activity of kinases such as PKA or GSK3β, which is a common feature of PKD pathogenesis (32–34), may increase GLI3 processing in the context of PKD. Previous studies have shown that overactivation of PKA activity within the context of PKD, leads to elevation of GLI3R and reduced GLI1 protein (34). These authors suggested that the enhanced GLI3R activity may contribute to the cystic phenotype, with mice exhibiting elevated GLI3R caused by constitutive PKA activation developing more severe PKD (34).

Following on from the identification of increased levels of *Gli3*, the effect of modulating the levels of *Gli3* was assessed in the *Cpk* mouse model of ARPKD and in a novel *in vitro* human model of ARPKD. Utilising a *Gli3* haploinsufficiency model, *Gli3* transcript levels were significantly reduced in *Cpk* mice, although this did not alter the cystic phenotype of *Cpk* mice. To further interrogate the function of *GLI3* in the pathogenesis of ARPKD, a human cellular model of ARPKD was then utilised. Initially, the Hh pathway was inhibited using the SMO-inhibitor cyclopamine, which reduced cyst size in control and *PKHD1*-mutant human collecting duct cells. Pharmacological inhibition of SMO reduces cystogenesis in other *in vitro* PKD models, including metanephric cystic explants and human primary ADPKD cells, as well as *in vivo* in the *Pck* rat model of ARPKD (8, 10, 12, 13). However, a specific reduction of *GLI3* transcript levels using siRNA did not alter cyst formation in these *PKHD1*-mutant HCD cells, indicating the increased *GLI3* within these collecting duct cells is not modulating cystogenesis.

The variation in *in vitro* cyst size observed after treatment with cyclopamine, compared with *GLI3* siRNA inhibition, can likely be attributed to cyclopamine’s more comprehensive inhibition of the Hh pathway. By antagonising SMO, cyclopamine reduces the level of GLI1 and GLI2 (35), which may be key to modulating cystogenesis in this model. Additionally, cyclopamine can have off target functions and has previously been demonstrated to inhibit growth and induce apoptosis, independently of SMO/GLI inhibition (36, 37). Thus, the changes observed in cyst size following cyclopamine treatment are likely due to the broader effect on the Hh pathway or off-target effects.

Although, pharmacological modulation of the Hh pathway has been demonstrated to reduce cystogenesis both *in vitro* in this study and *in vitro* and *in vivo* in previous reports (10, 12–14), there is limited data suggesting that the genetic modulation of this pathway alters PKD cystogenesis (10, 15). Prior studies have not investigated the modulation of *GLI3* specifically in cellular models of either ADPKD or ARPKD. Genetic manipulation of *Gli3* in epithelial cells has previously been analysed in the context of ADPKD. The loss of *Gli2* and *Gli3* did not alter cyst formation caused by mutations in *Pkd1*, indicating that the downstream Hh transcription factors, *Gli2* and *Gli3,* have no role within the tubular epithelium in mediating cystogenesis in ADPKD (15). This matches the findings in this report which demonstrated no difference in cyst formation and progression in ARPKD, both in a mouse model and in *PKHD1-*mutant human collecting duct cells, following Gli3 global repression.

A key limitation of this study is the level of GLI3 downregulation achieved with these models. It should be considered that complete abrogation of the gene might impact cyst formation and progression, however this was not feasible in our *in vivo* model. *Gli3^XtJ/XtJ^* mice are embryonic lethal and therefore it is not possible to analyse mice with a complete loss of *Gli3* expression postnatally (38). In our *in vitro* model, siRNA inhibition was able to reduce the levels of *GLI3* down to that observed in controls, although techniques like CRISPR-Cas9 may be of use to study the total loss of *GLI3* in this context. Additionally, as GLI3R protein levels were elevated in *Cpk* mice, another approach could be to study the effect of altering the level of GLI3R on cyst formation and progression in the context of ARPKD.

Lastly, this report describes the development of a human cellular model of ARPKD. A significant obstacle to advancing our understanding of the pathogenic mechanisms involved in ARPKD is the absence of genetically relevant models of ARPKD (39). In recent years, there has been progress in the development of cellular ARPKD models as a complementary approach to mouse models. *PKHD1^-/-^* ureteric epithelial cells isolated from human iPSC kidney organoids spontaneously form cyst-like structures when induced to form ureteric bud stalks (40). Additionally, two studies have described the use of organoids derived from ARPKD patient-derived iPSCs, although these did not reliably form cysts without cAMP stimulation. Moreover, in both studies these cystic tubules formed in LTL+ proximal tubules with only some developing in CDH1+ distal nephron regions (39, 41). CRISPR-Cas9 has also been employed in iPSCs to produce a *PKHD1-*mutant ARPKD kidney organoid-on-a-chip model. Organoids were exposed to fluid-flow on 3D millifluidic chips, resulting in the formation of cyst-like structures in distal nephrons only in *PKHD1^-/-^* organoids and not in *PKHD1^+/-^* organoids (39).

We have described a novel human *PKHD1*-mutant cellular cyst model, which can be utilised for the study of ARPKD and complement current ARPKD organoid models. This *PKHD1*-mutant cell line contains a compound heterozygous *PKHD1* mutation in a region of the gene which contains several pathogenic variants in patients (22). Compound heterozygous mutations occur in the majority of ARPKD cases (42, 43), thus this *PKHD1-*mutant line provides a valuable model of ARPKD. The *PKHD1*-mutant collecting duct cells form larger spherical cysts in 3D culture than isogenic controls over 6 days without the need of additional cAMP stimulation or fluid-flow. A key advantage of this model is the formation of collecting duct cyst-like structures, which is the primary cell type in which cysts form in patients with ARPKD (1). However, a limitation of the assay is that it lacks the cellular complexity and microenvironment that would be present in ARPKD tissue, such as the vasculature, immune or surrounding interstitial cells, which are important to fully mimic the cystic environment. Additionally, these *PKHD1*-mutant cysts are responsive to drug treatment quantified by cyst size, demonstrated using the drug cyclopamine. Hence, this cellular model is highly valuable due to its ease of use and scalability as well as having the potential to be utilised in applications such as high throughput drug screening, facilitating the evaluation of novel therapeutic interventions.

Overall, we have demonstrated that although levels of GLI3 are enhanced in models of ARPKD, dampening this upregulation does not alter cyst progression in both mouse and human models, highlighting the functional complexity of the Hh pathway in ARPKD. In addition, we have developed a cellular model of human ARPKD which provides a crucial resource for studying the pathogenesis of the disease, as well as discovering novel therapeutic targets and drugs aimed at addressing the needs of individuals with ARPKD.

## Acknowledgements

LGR was supported by a Kidney Research UK (KRUK) PhD studentship (ST_006_20181123) and a University College London Research Culture Award. This work was also supported by a Wellcome Trust Investigator Award (220895/Z/20/Z) and a BBSRC International Partnership Institutional Award (BB/X512011/1) to DAL, as well as a Canadian Institutes of Health Research Foundation Grant to NDR. The authors would like to thank Professor Pierre Ronco (Hopital Tenon, Paris, France) for providing the human collecting duct cell line.

## Conflict of interest statement

All authors declare no conflicts of interest.

## Author contributions

LGR, NDR, PJW and DAL conceived and designed the study. LGR and MKJ acquired mouse material. TC provided the *Gli3^XtJ^* mouse line. LGR performed mouse husbandry, qRT-PCR, western blotting, renal function assays and histology, with assistance from MKJ, LW, JCC and CJR. NPT, GS, KLP designed the guide RNAs and generated the CRISPR plasmids. LGR generated and characterised the *PKHD1*-mutant cell line and performed drug and siRNA treatments. LGR collated the data and presented the figures. LGR and DAL wrote the manuscript, and subsequently all authors were involved in revision and preparation of the final manuscript.

---

**Supplementary Figure 1.**
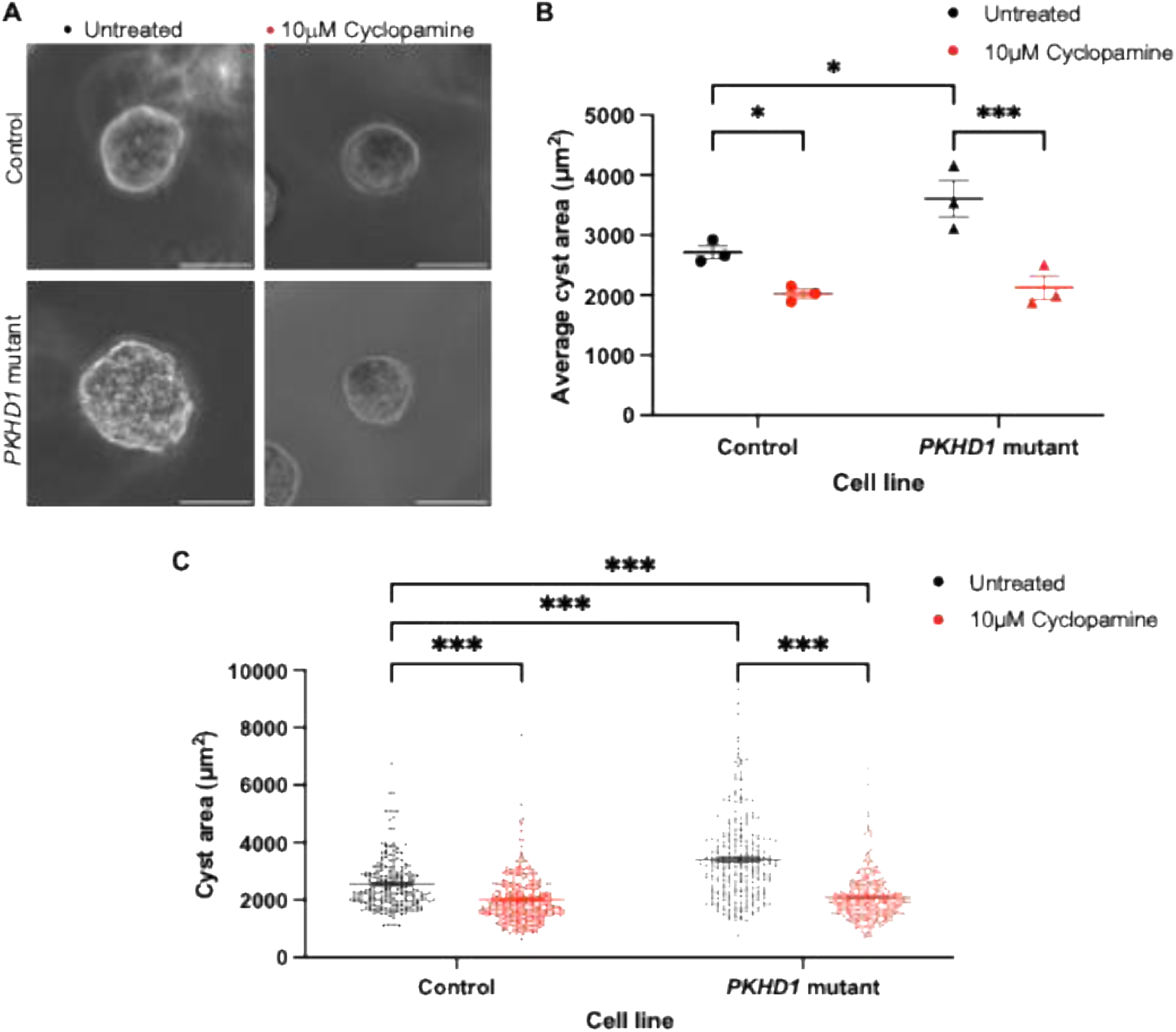
Cyclopamine treatment inhibits ARPKD cyst formation *in vitro*. Isogenic wildtype control and *PKHD1*-mutant human collecting duct cells were embedded in Matrigel and cultured for 6 days to generate 3D cyst-like structures. Cells were left untreated or treated with 10μM Cyclopamine, a SMO inhibitor, every 2 days. (**A**) Representative images of isogenic wildtype control and *PKHD1*-mutant 3D cyst-like structures at day 6, following 10μM Cyclopamine treatment. (**B**) The cross-sectional area of each cyst was quantified, and the average cyst area was calculated for each independent repeat. (**C**) Cyst area of each individual cyst quantified across all repeats. n=3 independent repeats. Data represents mean ± SEM. Two-way ANOVA with Tukey’s multiple comparisons test. Statistical comparisons are shown relative to the untreated isogenic wildtype control and between conditions of the same cell type. *P<0.05, ***P<0.001.

## Notes

### Competing Interest Statement

The authors have declared no competing interest.

